# Together Inbreeding and Reproductive Compensation Favor Lethal *t*-Haplotypes

**DOI:** 10.1101/2023.07.26.550691

**Authors:** Manisha Munasinghe, Yaniv Brandvain

## Abstract

Male mice who are heterozygous for distorting and non-distorting alleles at the *t*-haplotype transmit the driving *t*-haplotype around 90% of the time – a drastic departure from Mendelian expectations. This selfish act comes at a cost. The mechanism underlying transmission distortion in this system causes severe sterility in males homozygous for the drive alleles, ultimately preventing its fixation. Curiously, many driving *t*-haplotypes also induce embryonic lethality in both sexes when homozygous, however, this is neither universal nor a necessity for this distortion mechanism. Charlesworth provided an adaptive explanation for the evolution of lethal *t*-haplotypes in a population segregating for distorting and non-distorting *t* alleles – namely, if mothers compensate by replacing dead embryos with new offspring (or by transferring energy to surviving offspring), a recessive lethal can be favored because it effectively allows mothers the opportunity to trade in infertile males for potentially fertile offspring. However, this model requires near complete reproductive compensation for the invasion of the lethal *t*-haplotype and produces an equilibrium frequency of lethal drivers well below what is observed in nature. We show that even low levels of systemic inbreeding, which we model as brother-sister mating, allows lethal *t*-haplotypes to invade with much lower levels of reproductive compensation. Furthermore, inbreeding allows these lethal haplotypes to largely displace the ancestral male-sterile haplotypes. Our results show that together inbreeding and reproductive compensation move expected equilibria closer to observed haplotype frequencies in natural populations and occur under more reasonable parameter combinations.

## Introduction

Mendel’s Law of Segregation dictates that, during meiosis, only one of two gene copies present in a diploid organism will be distributed to each gamete with equal probability for each allele (Abbott and Fairbanks 2016). At most loci, alternative alleles are fairly transmitted – abiding by Mendel’s Law of Segregation; however, there are selfish genetic elements that are not only capable of biasing this equal distribution in their favor but are also quite effective in doing so. Segregation distorters are one such class of selfish genetic elements. As the name suggests, segregation distorters are capable of influencing meiosis in order to distort allelic segregation such that they are transmitted over 50% of the time to offspring. Segregation distorters have been discovered in several distinct populations and species, but the *t*-haplotype system in house mouse and related species of *Mus* is a seminal example (another being the *SD* gene complex found in natural populations of *Drosophila melanogaster*) (Zimmering et al. 1970; Lyttle 1991; Silver 1993; Larracuente and Presgraves 2012; Lindholm et al. 2016).

The mouse *t*-haplotype itself is a large variant (30 - 40Mb) on chromosome 17 that contains hundreds of genes and several large inversions. These inversions minimize recombination between the *t* and wild type haplotypes (Burt and Trivers 2006; Herrmann and Bauer 2012; Kelemen and Vicoso 2018). Housed within the *t*-haplotype are a series of genes that work in tandem to render wild-type sperm non-functional (Artzt et al. 1982; Herrmann et al. 1986; Olds-Clarke and Johnson 1993; see Schimenti 2000 for a more thorough explanation of the molecular mechanisms underlying *t*-haplotype transmission). Consequently, the *t*-haplotype segregates normally in females but distorts meiosis (i.e., is transmitted to offspring > 50% of the time) in males. This gametic distortion is extreme – in a single mating by heterozygous males, the *t*-haplotype is transmitted around 90% of the time (Silver 1993; Lyon 2003). However, as a consequence of the underlying drive mechanism, males homozygous for the *t*-haplotype are rendered sterile (Lyon 1984). Further adding to the complexity of the *t*-haplotype system is the presence of recessive lethal alleles that are strongly linked to the haplotype. Of the 16 well characterized *t*-haplotypes, 13 of them carry a recessive lethal mutation (Klein et al. 1984). Individuals homozygous for the lethal allele die early during embryogenesis, regardless of their chromosomal sex, but the exact time of action varies between different lethal haplotypes (Bennett 1975).

*t*-haplotypes are found throughout the *Mus musculus* species complex, which includes *M.m. musculus*, *M.m. domesticus*, *M.m. castaneus*, and *M.m. bactrianus* (Silver 1993; Huang et al. 2001; Dod et al. 2003). Investigations into the frequency of *t*-haplotypes within wild populations have given variable results. Before continuing, it is important to note that house mouse populations are often demic in structure (Klein and Bailey 1971). This means a single population often consists of several smaller breeding groups that are predominantly isolated with low levels of migration between distinct demes. Within a population, estimates for the frequency of *t*-haplotypes, which has historically been determined based on the number of heterozygous individuals, average between 13 -25% (Dunn et al. 1960; Anderson 1964; Petras 1967; Klein et al. 1984; Lenington et al. 1988; Ruvinsky et al. 1991); however, this frequency can vary substantially across demes and years sampled for a given population. Petras (1967) characterized the frequency of *t*-haplotypes in wild populations at five localities from various buildings over a four year period and found the overall frequency to be approximately 16% (Petras 1967). In a given locality, one building could entirely consist of mice heterozygous for the *t*-haplotype while another building could lack it entirely. Tracking pooled estimates across buildings for a given locality shows that the frequency of the *t*-haplotype fluctuates over time (in one locality the frequency changed from ∼0.308 to 0.026 in a single year). Similarly, Ardlie and Silver (1998) performed a large survey of *M.m. domesticus* and found the frequency of the *t*-haplotype to vary between 0 - 70% across independent populations (Ardlie and Silver 1998). The overall frequency for the *t*-haplotype was only 0.062, due to the absence of the *t*-haplotype in approximately half of the populations sampled.

The low and variable observed frequency of the *t*-haplotype conflicts with deterministic population genetic theory on the evolution of the *t*-haplotype. The effect of the *t*-haplotype was originally thought to be caused by a single gene. Consequently, early models often considered the evolution of two alleles, either wild-type or *t* where the *t*-allele is either lethal or male-sterile. These models show that both the lethal and male-sterile *t*-allele can invade and persist at a stable equilibrium within populations; however, these expected equilibrium frequencies are often much higher than observed values (Bruck 1957; Dunn and Levene 1961). For example, Dunn and Levene 1961 showed that, with a segregation distortion of 0.85, a male-sterile *t*-allele should reach an equilibrium of 0.7. Empirical results in their study on a sampled wild population harboring a male-sterile *t*-haplotype show an observed frequency of 0.37, just over half as small as the expected frequency (Dunn and Levene 1961). Interdemic selection – i.e. local extinction of demes in which the t-haplotype is common – between fully isolated demes of small size can generates both stochastic heterogeneity in the frequency of the *t*-haplotype across demes, and a lower overall population frequencies for the *t*-haplotype (Lewontin and Dunn 1960; Lewontin 1962; Levin et al. 1969). However, models that extend this to incorporate even a small amount of migration (1%) between demes rarely predict natural *t*-haplotype frequencies under biologically plausible parameter combinations (Levin et al. 1969; Durand et al. 1997). Consequently, interdemic selection on its own cannot sufficiently explain *t*-haplotype evolution.

Two intriguing factors that have been considered independently for *t*-haplotype evolution are inbreeding and reproductive compensation. Inbreeding is pervasive among natural populations of *Mus musculus*, but it can vary substantially within and between populations (Yamazaki et al. 1976; Hurst et al. 2001; Sherborne et al. 2007; Morgan et al. 2022). Early models considering interdemic selection informally explored inbreeding, as small deme size functionally results in inbreeding; however, as mentioned above, models of interdemic selection fail to predict observed *t*-haplotype frequencies. Petras 1967 incorporated a coefficient of inbreeding, that can be considered as a measure of the Wahlund effect (i.e., the numerical reduction of heterozygotes due to population subdivision), into a two-allelic model of *t*-haplotype evolution and found stable equilibria that were much closer to observed estimates (Petras 1967). These models, however, do not consider the social interaction of brother-sister mating that formally couples the reproductive success of relatives, which is expected to further limit the spread of *t*-haplotypes.

Reproductive compensation, on the other hand, assumes that early embryonic lethality may serve an adaptive purpose. Early embryonic elimination of recessive lethals may improve the postnatal survival of remaining offspring, increase the duration of reproductive life for mothers by reducing litter size, or simply have minimal effect on female fecundity because more embryos are implanted than come to term (Batten and Berry 1967; Priestnall 1972; Roberts 1981). Charlesworth 1994 formally considered the kin selection advantage to early embryonic lethality via reproductive compensation. This model demonstrates that reproductive compensation can facilitate the invasion of a lethal haplotype into a population jointly polymorphic for a wild-type and male-sterile haplotype (Charlesworth 1994). However, this requires highly favorable invasion parameters (i.e., the *t*-haplotype has no effect in heterozygotes and is fully male-sterile), strong segregation distortion, and high values of reproductive compensation to maintain the lethal *t*-haplotype at substantial frequencies (> 20%). There were no parameter combinations where the lethal successfully displaced the male-sterile haplotype, which conflicts with empirical observations as the lethal *t*-haplotype is often found at higher frequencies than the male-sterile. Recent evidence suggests that the impact of reproductive compensation is likely much lower, if present at all, than the values needed in Charlesworth 1994 to even get invasion of the lethal *t*-haplotype (Lenington et al. 1994; Lindholm et al. 2016).

Here, we consider the influence of reproductive compensation and inbreeding via sib-mating on *t*-haplotype evolution. We start by looking at a two-allelic model to explore when a male-sterile *t*-haplotype can invade a naive population and how inbreeding impacts the final equilibrium frequency. We then transition to a three-allelic model with the wild type, male-sterile *t*-haplotype, and lethal *t*-haplotype. We first introduce the male-sterile *t*-haplotype and, if a stable polymorphism is achieved, consequently introduce a lethal *t*-haplotype. Across these models, we explore how reproductive compensation and inbreeding interact and influence the invasion dynamics for the lethal *t*-haplotype. We assess whether these forces in conjunction result in stable equilibrium frequencies closer to those observed in natural populations, as previous models have struggled to match empirical observations. We also examine under which conditions the lethal *t*-haplotype displaces, or reaches higher frequency than, the male-sterile haplotype. We compare our results to both the Petras 1967 and Charlesworth 1994 model to explore how reproductive compensation and inbreeding both independently and jointly influence *t*-haplotype evolution.

## Methods

We leverage the framework first established in Charlesworth 1994 and extend it to consider inbreeding via sibling-mating. We consider a single population of infinite size with at most three haplotypes at the *t*-complex - (1) wild type (*A*), (2) the male-sterile *t*-haplotype (*a*), and the lethal *t*-haplotype (*aL*). We will often refer to the male-sterile *t*-haplotype as the distorter haplotype, the lethal *t*-haplotype as the lethal distorter haplotype, and the *t*-haplotype as the combination of the distorter and lethal distorter haplotype. The distorter haplotype (*a*) has no effect in females, but males homozygous for the distorter (*aa*) experience a reduction in fertility of 1 - *t* relative to wild-type homozygotes. It is possible to parameterize a reduction in fitness for heterozygotes (*s* denotes a reduction in fitness of 1 - *s* in heterozygotes relative to wild-type homozygotes); however, this impedes the invasion of the *t*-haplotype. Consequently, we assume there is no effect (*s* = 0) in heterozygotes to limit the number of parameters explored to improve interpretability of our results. Heterozygous males will, however, transmit the distorter allele to their offspring with probability *k*. The lethal distorter (*aL*) haplotype shares many of the same features as the distorter haplotype. The key difference is that all individuals homozygous for the lethal distorter are assumed to be killed early during embryogenesis. Charlesworth 1994’s model technically considered the lethal distorter haplotype as two distinct alleles and parameterized recombination between them; however, as his results show that extremely tight linkage is required for the lethal to invade, we treat this as a single allele (*aL*) mimicking the case where those two alleles are in complete linkage (*r* = 0) with each other. This allows us to consider this framework as a three-allele model instead of as a two-locus, two-allele model.

Charlesworth 1994’s initial model considered the influence of reproductive compensation, which occurs as a result of early termination of embryos homozygous for the lethal distorter. Consequently, the net reproductive success of mothers is not reduced in proportion to the frequency of lethals in their litter. Charlesworth 1994 models this with the parameter *C*, which assigns the degree of reproductive compensation (*C* = 1 implies the mortality of lethals has no effect on female fitness). If the frequency of lethal offspring from a given mating type is 𝑝_𝑚_, then the expected number of viable offspring produced by this mating is adjusted by (1 − 𝑝_𝑚_)/(1 − 𝑝_𝑚_)^𝐶^. We replicate this model and affirm Charlesworth 1994’s results. Even with highly favorable invasion parameters (*s* = 0, *t* = 1) and a strong segregation distortion (*k* = 0.9, 0.99), the lethal distorter is maintained at relatively low frequencies and fails to displace the distorter allele (Fig S1).

In our model, we parameterize inbreeding as the probability that a female mates with her brother. That is, with probability *psib*, a female mates with her brother, and with probability *1 - psib*, a female chooses a random mate from the population. Implementing this model requires us to know the probability of a brother’s genotype given his sister’s genotype. Representing brother genotype as *Bi*, and sister genotype as *Sj*, we must find *p*(*Bi* | *Sj*). To solve this, we first find the probability that a sister of genotype *j* was born into family type *k* (i.e. *p*(*Fk* | *Sj*)). Using Bayes’ theorem, we can define this as follows in Equation 1.

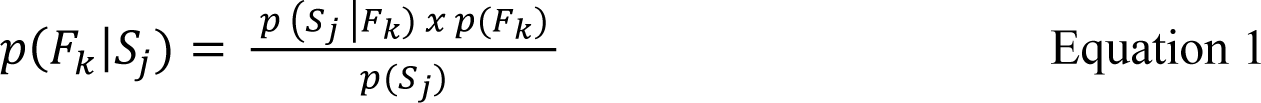

Values on the right hand reflect frequencies calculated after considering transmission distortion and selection within families in the previous generation. Consequently, *p*(*Fk*) is the frequency of family *k* after viability and fertility selection in the previous generation (i.e., 𝑝(𝐹_𝑘_) = (𝑝(𝐹_𝑘(𝑡−1)_) × 𝑤̅_𝑘_)/𝑤̅). 𝑝(𝑆_𝑗_|𝐹_𝑘_) is similarly the proportion of sisters (or daughters) of genotype *j* generated from family type F*k*, while 𝑝(𝑆_𝑗_) is the proportion of daughters of that genotype (both after selection).

We can then use the law of total probability to find *p*(*Bi* | *Sj*). This simply becomes Equation 2, which is the product of the probability that a sister of genotype *j* was born into family type *k* and the probability that a brother has genotype *i* given he was also from family *k* (i.e. 𝑝(𝐵_𝑖_|𝐹_𝑘_)), summed over all families.

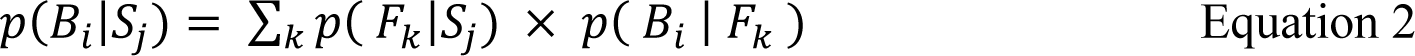

Overall, we consider four distinct models. Models 1 and 2 represent two-allele models where we can track the evolution of a male-sterile *t*-haplotype into a naive population with and without the probability of brother-sister mating. Models 3 and 4 extend this framework to consider a three-allele model where we first introduce a male-sterile *t*-haplotype into a naive population and then, if a stable polymorphism can be obtained for the distorter, we introduce a lethal *t*-haplotype. There are four key parameters across our models: *t, C, k,* and *psib* (Table 1). *t* represents the reduction in fitness in homozygotes. As a reminder, the male-sterile *t*-haplotype (*a*) only has fitness reductions in males, while the lethal *t*-haplotype (*aL*) reduces fitness in both sexes. *C* represents the degree of reproductive compensation induced by the embryonic lethality of the *t*-haplotype.

**Table 1.** Key Parameters Across Our Models. All parameters explored in our models. Column 1 shows the variable name, column 2 details what the parameter represents, column 3 shows the explored parameter values, and column 4 shows which models that parameter is present in. Model 1 and 2 represent our two-allele models, while Models 3 and 4 represent our three-allele models.

| Variable | Parameter Value | Tested Values | Models Present |
| --- | --- | --- | --- |
| $t$ | Fitness Effect in Homozygotes | [0, 0.1, 0.2, 0.3, 0.4, 0.5, 0.6, 0.7, 0.8, 0.9, 1.0] | 1 - 4 |
| $C$ | Reproductive Compensation | [0, 0.1, 0.2, 0.3, 0.4, 0.5, 0.6, 0.7, 0.8, 0.9, 1.0] | 3, 4 |
| $k$ | Segregation Distortion | [0.5, 0.6, 0.7, 0.8, 0.9, 0.99, 1.0] | 1 - 4 |
| $p_{sib}$ | Proportion of Sib-Mating | [0, 1e-4, 1e-3, 0.01, 0.025, 0.050, 0.075, 0.1, 0.2, 0.4, 0.5] | 2, 4 |

All models begin by introducing the male-sterile *t*-haplotype into a naive population. We always introduce the male-sterile *t*-haplotype by adjusting the frequency of *Aa* heterozygotes to 1/1000 and AA homozygotes to 1 - 1/1000, which equates to a starting allele frequency of 1/2000 for the male-sterile haplotype. We move forward in generational time until the change in allele frequency for the male-sterile *t*-haplotype is less than 1 x 10^-4^ after at least 100 generations. For the lethal distorter, its starting frequency depends on the equilibrium point of the male-sterile haplotype. If the proportion of *Aa* heterozygotes is greater than 0.05, we reduce the proportion of *Aa* heterozygotes by 0.05 and increase the proportion of *AaL* heterozygotes from 0 to 0.05. If not, we simply divide the proportion of *Aa* heterozygotes by 2 and increase the proportion of *AaL* heterozygotes by an equal amount. We do this to minimize the amount of the parameter space where the introduction of the lethal *t*-haplotype (*aL*) is less than 0.05, as we found that small starting allele frequencies for the lethal often took very long to reach an equilibrium point due to very small increases in allele frequency during earlier generations. We once again move forward in generational time until the change in allele frequency for the lethal *t*-haplotype is either less than 1 x 10^-8^ or less than 0.005 in frequency after at least 100 generations. We use a frequency less than 0.005 as a proxy for loss as allele frequencies were slowly tending towards 0.

Models with inbreeding (2 and 4) decrease the proportion of random mating by parameter *psib* and replace the remaining proportion with mating generated via full sib-mating. The overall frequency of each mating type is then the proportion of that mating type generated by random mating plus the proportion generated by sib-mating (Table 2a). We then establish gametic frequencies in both males and females. In females, alleles segregate normally. In heterozygous males, however, the distorter allele (either *a* or *aL*) is transmitted to offspring with frequency *k*.

**Table 2.**
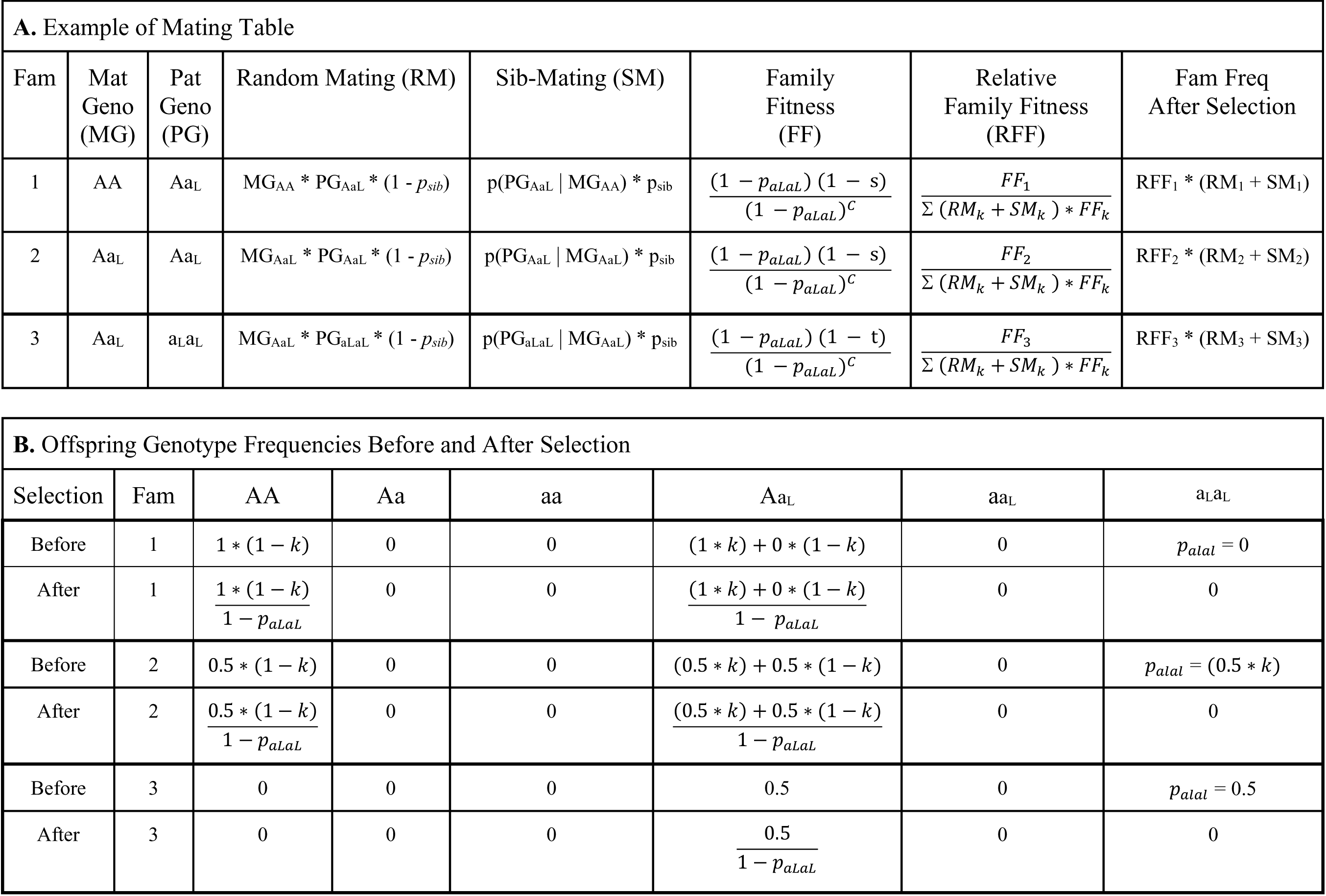
Mating Table and Offspring Genotype Frequencies for 3 Example Mating Types for Model 4. A. Example of Mating Table **B.** Offspring Genotype Frequencies Before and After Selection. Model 4 is our three-allele model which includes a lethal *t*-haplotype with reproductive compensation and a proportion of inbreeding via sib-mating.

From these gamete frequencies, we can establish the genotype frequencies among generated offspring which we then adjust based on family fitness (Table 2b). In models without reproductive compensation, family fitness depends solely on the male genotype (with homozygous males experiencing a reduction in fitness equal to *t*). Models with reproductive compensation not only adjust family fitness based on the male genotype but also by the frequency of lethal offspring generated adjusted by the coefficient of compensation (i.e., (1 − 𝑝_𝑚_)/(1 − 𝑝_𝑚_)^𝐶^. where 𝑝_𝑚_ is the frequency of lethal offspring generated from a given mating type and *C* is the extent of reproductive compensation). Additionally, we consider special cases of Models 3 and 4 where we introduce only the lethal distorter, which are functionally analogous to a two-allele model with only the wild type and lethal distorter haplotypes.

Across our four models, we can determine when both the male-sterile and lethal *t*-haplotype invade and reach stable polymorphic equilibrium. We can also explore when the lethal *t*-haplotype is capable of displacing the male-sterile haplotype (i.e., the lethal *t*-haplotype surpasses the male-sterile in frequency), as the lethal *t*-haplotype is often found at higher frequencies than the male-sterile in natural populations. Ultimately, we can assess how reproductive compensation and inbreeding via sib-mating influence *t*-haplotype evolution and whether their interaction moves predicted allele frequencies closer to observed estimates.

## Results

### Inbreeding decreases the frequency of the t-haplotype

The current hypothesis for the evolution of the *t*-haplotype assumes that the distorter allele evolved first, such that homozygous males were sterile but viable, followed by the evolution of recessive lethal mutation(s) on the distorter haplotype. We first evaluated how inbreeding influenced the invasion trajectories for the male-sterile *t*-haplotype and whether stable polymorphic equilibrium could be reached. We find that the distorter allele can successfully invade and reach stable polymorphic equilibria under the explored parameters for inbreeding; however, as the probability of sib-mating increases, the final frequency of the distorter allele is reduced (Fig 1a). Subtle amounts of sib-mating have negligible effects on the final frequency of the distorter haplotype with at least a 1% chance of sib-mating required to reduce the allele frequency by at least 1%. For example, an effective distorter (*k* = 0.9) with complete male sterility (*t* = 1), a 10% chance of sib-mating reduces the stable frequency of the distorter allele by ∼7.5% relative to no sib-mating (0.74 relative to 0.8 respectively) with even more severe reductions as the chance of sib-mating increases. Unsurprisingly, for a fixed probability of sib-mating, the final frequency of the distorter allele increases (and can fix) as the fitness effect in homozygous males (*t*) decreases and as the strength of the segregation distortion (*k*) increases (Fig 1b). In sum, lower fitness effects in homozygous males, stronger segregation distortion, and reduced levels of inbreeding all facilitate the invasion of a male-sterile *t*-haplotype.

**Figure 1.**
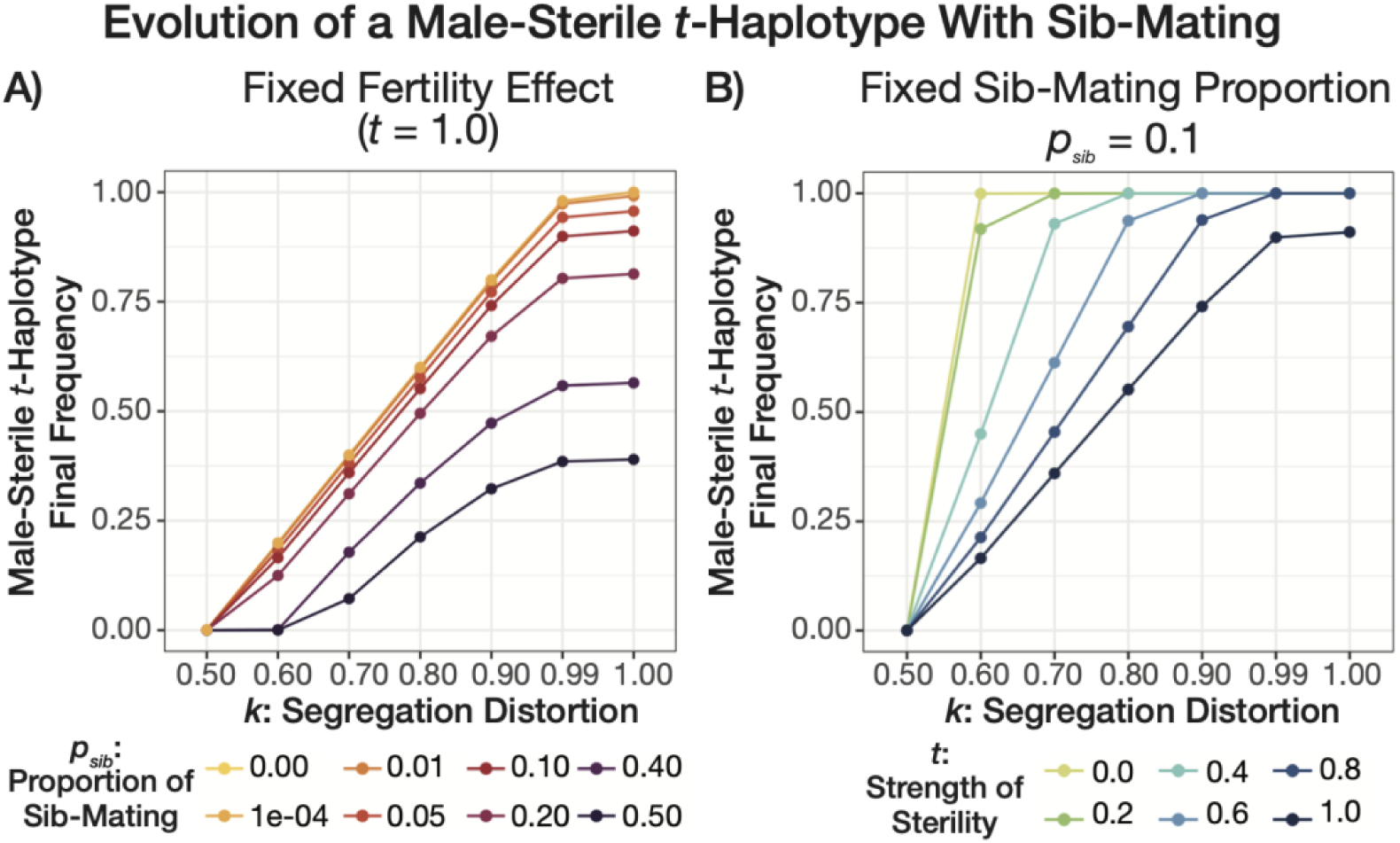
Allele Frequency Equilibria for Male-Sterile *t*-Haplotype With Sib-Mating. Model 2 considers the invasion criteria for a male-sterile *t*-haplotype into a naive population. The x-axis of both plots shows the strength of the segregation distortion (*k*), while the y-axis show the final allele frequency for the male-sterile *t*-haplotype. (A) We fix the fitness effect in male homozygotes (*t* = 1.0) and plot the final allele frequencies by the proportion of sib-mating (represented by the differently colored lines ranging from light yellow to dark purple with darker colored lines representing more sib-mating). (B) We fix the proportion of sib-mating (*psib* = 0.1) and now plot the final allele frequencies by the fitness effect in male homozygotes (represented by differently colored lines ranging from light green to dark blue with darker colored lines representing a stronger deleterious effect in homozygous males).

We did consider whether a lethal distorter could invade relative to the wild type haplotype. We did this by exploring a special case of our three-allele models where we set the frequency of the male-sterile distorter to zero and only introduced the lethal distorter. Under random mating, our results perfectly align with analytical equilibrium values predicted by Dunn and Levene 1961 for a lethal *t*-haplotype (Fig S2 - note, these analytical equilibria assume *s* = 0 and *t* = 1). Petras 1967 is the only alternative model to incorporate inbreeding into models of *t*-haplotype evolution (via deviations from Hardy-Weinberg equilibrium due to population structure). Under low proportions of sib-mating (< 10%) our results are roughly comparable to expected frequencies derived from Petras 1967. Similarly to the male-sterile *t*-haplotype, inbreeding via-sib mating does allow the lethal *t*-haplotype to invade but results in lower equilibrium values. However, once the proportion of sib-mating increases, the Petras 1967 expected frequencies deviate strongly from both our frequencies and observed expectations (Fig S3). In contrast, our model which directly links the reproductive success of relatives stays within much more reasonable final frequencies under higher levels of inbreeding (ranges from 0.09 to 0.33).

### Together inbreeding and reproductive compensation favor lethal t-haplotypes

We then built on Charlesworth’s (1994) model and asked whether inbreeding facilitated the invasion of a lethal drive allele in a population segregating for the wild-type and the male-sterile *t*-haplotype. Earlier we demonstrated that regular inbreeding via sib-mating reduces the equilibrium frequency of a male-sterile distorter. Here we find that even rare sib-mating allows the lethal *t*-haplotype to invade the population under much lower values of reproductive compensation than observed in the random mating model. For example, under favorable parameters (*s* = 0, *t* = 1, and *k* = 0.99), reproductive compensation (*C*) needed to be greater than 0.85 in order for the lethal *t*-haplotype to invade a randomly mating population (Fig S2 and Charlesworth 1994). We find that, under similar conditions, the lethal *t*-haplotype can not only invade under lower reproductive compensation values but also fully displace the male-sterile haplotype with sufficient levels of inbreeding (Fig 2). If we fix *C* at 0.6, which is a much more modest and potentially more reasonable estimate for reproductive compensation, we find that with a 1% chance of sib-mating the lethal *t*-haplotype can invade and reach an equilibrium frequency of 0.09. However, in this case, all three alleles are maintained with the male-sterile segregating at the highest frequency (∼ 88%). Slightly more inbreeding allows the lethal t-haplotype to further displace the male-sterile *t*-haplotype. For example, with a 5% probability of sib-mating under the same remaining parameters (*s* = 0, *t* = 1, *k* = 0.99, *C* = 0.6), the lethal *t*-haplotype nearly eliminates the male-sterile haplotype – reaching an allele frequency of 0.49. As a reminder, a frequency of 0.5 for the lethal *t*-haplotype represents fixation as homozygotes die during embryogenesis. If we decrease the strength of distortion (*k*), we find that higher sib-mating proportions were required for the lethal *t*-haplotype to invade and displace the male-sterile haplotype (Fig 2B).

**Figure 2.**
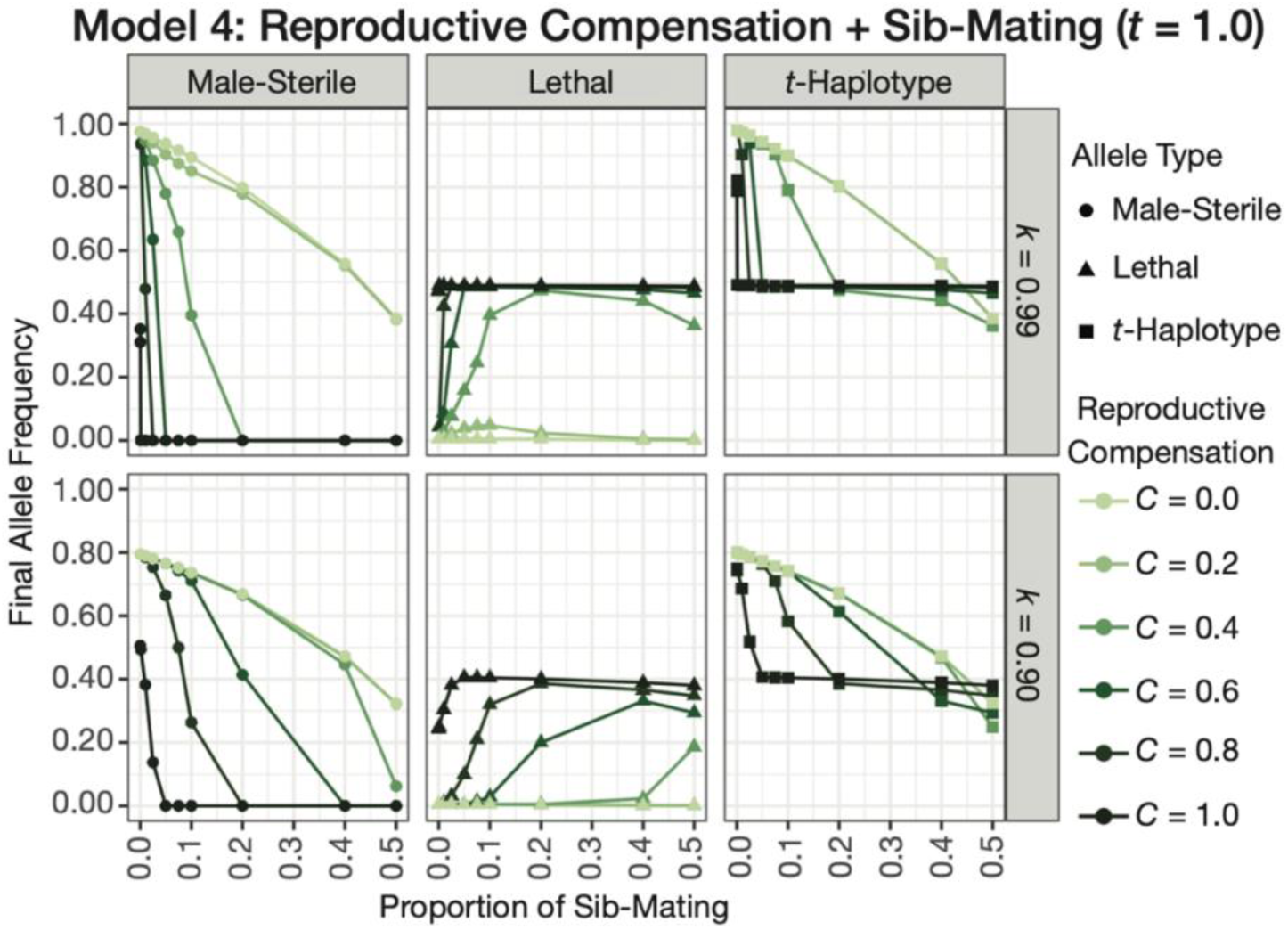
Allele Frequency Equilibria for Male-Sterile and Lethal *t*-Haplotypes With Sib-Mating. Model 4 considers the invasion criteria for a lethal *t*-haplotype into a population that is stably polymorphic for both the wild type and male-sterile *t*-haplotype. The x-axis of each plot shows the proportion of sib-mating, and the y-axis shows the final allele frequency. Each column shows the final allele frequency for either the male-sterile *t*-haplotype (left), lethal *t*-haplotype (middle), or *t*-haplotype (right). The *t*-haplotype frequency is simply the sum of the male-sterile and lethal *t*-haplotype frequencies. The first row shows results for a segregation distortion *k* = 0.99, while the second row shows this for *k* = 0.90. Points on each line show the observed frequencies with shape once again highlighting which allele we are investigating. The color of each line represents the extent of reproductive compensation (*C*) with darker colors of green indicating higher degrees of reproductive compensation.

Displacement of the male-sterile allele occurs under very low levels of sib-mating once reproductive compensation becomes sufficiently high. Once again, assuming the same favorable parameters, the lethal *t*-haplotype functionally displaces the male-sterile *t*-haplotype when *C* = 0.8 and *psib* = 0.025 (the final allele frequencies are 0.49 and 2.0e-05 respectively). By extending Charlesworth 1994’s model to allow for inbreeding which explicitly links the reproductive success of relatives, our model is capable of finding more reasonable parameter combinations under which the lethal *t*-haplotype can not only invade populations but also displace the male-sterile *t*-haplotype (Fig 3). At least some reproductive compensation (C > 0.4) is generally required for the lethal *t*-haplotype to invade which, while more reasonable, may still be higher than values estimated from natural populations. It is also worth noting that there may be a limit to the effectiveness of inbreeding. We did observe very subtle reductions (∼ 0.01) in the final allele frequencies reached under the highest proportions of sib-mating we explored.

**Figure 3.**
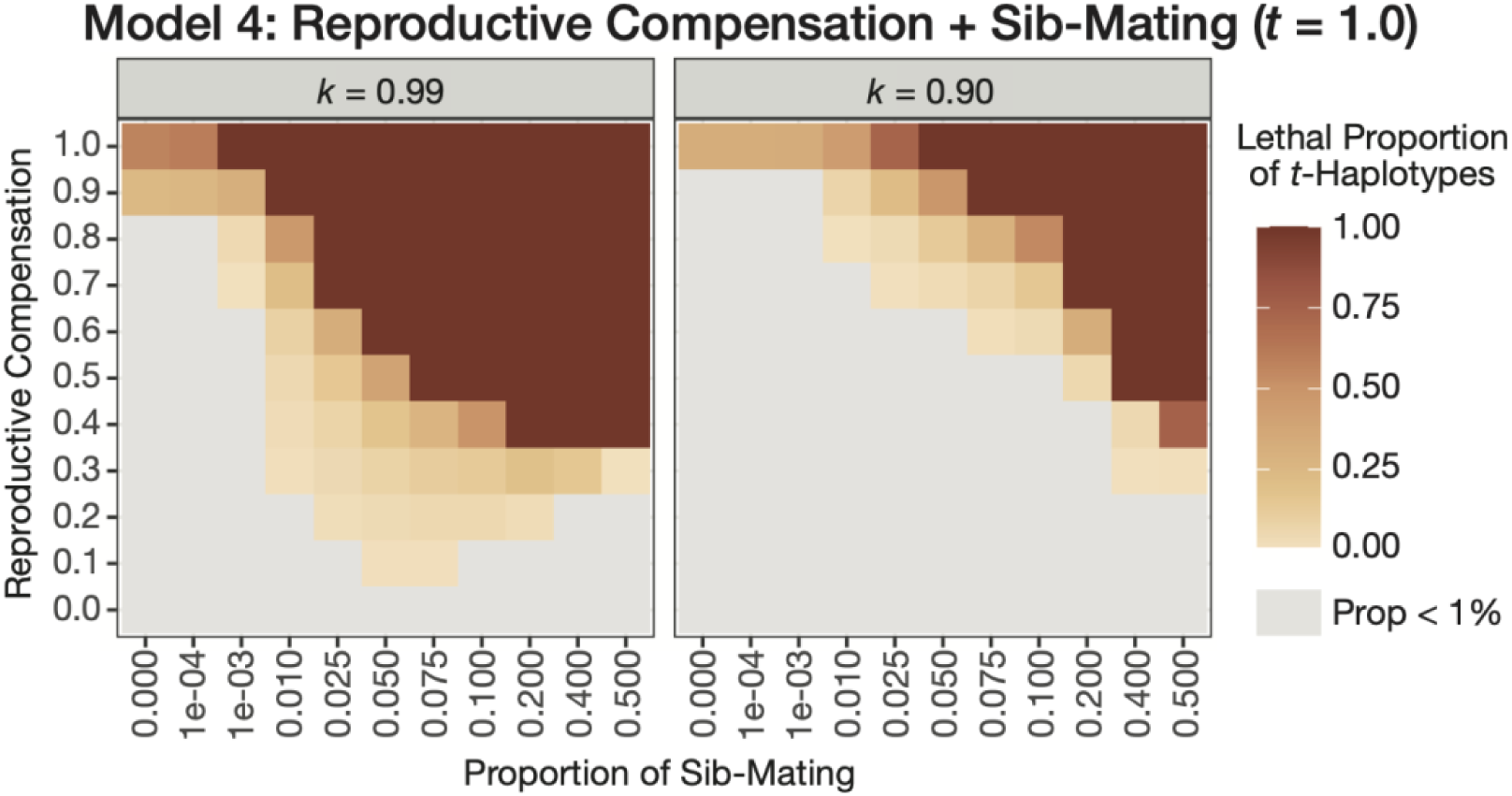
Final Lethal *t*-Haplotype Proportion For Model 4 with Reproductive Compensation and Inbreeding. Model 4 considers the invasion criteria for a lethal *t*-haplotype into a population that is stably polymorphic for both the wild type and male-sterile *t*-haplotype. The x-axis of each plot shows the proportion of sib-mating, and the y-axis shows the amount of reproductive compensation. Each cell shows the final proportion of present lethal *t*-haplotypes of the total *t*-haplotypes. The first column shows results for a segregation distortion *k* = 0.99, while the second row shows this for *k* = 0.90. The color each cell shows the overall proportion with darker colors of brown indicating higher proportion of lethal *t*-haplotypes. Grey cells indicate parameter combinations where the proportion of lethal *t*-haplotypes were less than 1%.

In sum, we find that, while neither inbreeding nor reproductive compensation on their own strongly favor the replacement of a male-sterile t-haplotype by a lethal t-haplotype, when combined these forces facilitate the invasion of a lethal *t*-haplotype. Removing a sterile male not only allows for its replacement by a fitter sibling (as in Charlesworth 1994) but also increases the reproductive success of his sisters by removing a sterile mate from the reproductive pool. Our models not only align with the results of previous theoretical work but also extend them to show how inbreeding and reproductive compensation in tandem favor the evolution of a lethal *t*-haplotype.

## Discussion

Charlesworth (1994) first demonstrated that reproductive compensation could provide an adaptive explanation for the presence of lethal *t*-haplotypes. While Charlesworth’s panmictic model showed that reproductive compensation could favor a lethal *t*-haplotype, it could only do so when females were near fully compensated for their lost embryos (C > 0.85), and this lethal driver could not reach appreciable frequencies. We show that introducing inbreeding to Charlesworth’s reproductive compensation model allows a lethal *t*-haplotype to invade when reproductive compensation is much less effective than required under panmixia. Additionally, inbreeding allows the lethal *t*-haplotype to become an appreciable portion of the present *t*-haplotypes, which aligns with natural observations.

Strikingly, the interaction between reproductive compensation and inbreeding is crucial for the displacement of a male-sterile *t*-haplotype by a lethal haplotype. Across all of our two-allele frameworks, which consider the evolution of a *t*-haplotype (either male-sterile or lethal) into a naive population, we find that inbreeding via sib-mating still allows for the invasion of the *t*-haplotype and results in lower equilibrium values than previous theory. This aligns with general theory that inbreeding reduces genetic conflict (Burt and Trivers 1998; Brandvain and Haig 2005; Bull 2016) and previous theory on the evolution of the *t*-haplotype in structured populations (Lewontin and Dunn 1960; Lewontin 1962; Petras 1967). While on its own, inbreeding does not favor the displacement of male-sterile *t*-haplotypes by lethal *t*-haplotypes, it does favor this displacement when embryo death is compensated. It does so by linking male sterility to the fitness of their sisters. Sisters of males homozygous for a male-sterile *t*-haplotype are stuck mating with an infertile brother, which decreases their inclusive fitness. In contrast, lethal *t*-haplotypes prevent such unsuccessful matings by eliminating these brothers before they can impact their sister’s fitness. This type of indirect selection also been shown to facilitate selection on maternally inherited genomes carrying male-harming mutations (Unckless and Herren 2009; Wade and Brandvain 2009), and it may explain the theoretical observation that selfing can – in some cases – hasten the spread of Medea-like distorters in Caenorhabditis. been previously shown to facilitate selection on male fertility on maternally inherited genomes (Noble et al. 2021).

We show that the interaction between reproductive compensation and inbreeding facilitates the adaptive replacement of male-sterile *t*-haplotypes by lethal *t*-haplotypes. However, questions remain as to whether this theoretical possibility actually aligns with and explains patterns in nature. There are many different lethal *t*-haplotypes which exhibit extreme variation in the timing of lethality. The exact time of lethality likely effects the extent of reproductive compensation associated with each such lethal haplotype (Bennett 1975). One lethal haplotype is expressed very early in development and is only associated with a 5% reduction in litter size (relative to observations for a similar cross with complementing *t*-haplotypes) suggesting the potential for extreme reproductive compensation, while other *t*-haplotypes in this same study suggested much more limited reproductive compensation (Lenington et al. 1994). A more recent survey on an Australian *t*-haplotype variant found no reduction in heterozygous female litter size when fertilized by heterozygous males, which suggests strong reproductive compensation for this variant (Manser et al. 2017). Conversely, a study of a different lethal *t*-haplotype found no evidence of reproductive compensation for the explored *t*-haplotype (Lindholm et al. 2013). Thus, while there is clearly an opportunity for reproductive compensation to influence the evolution of some *t*-haplotypes, it seems that many lethal haplotypes exist with limited opportunity for reproductive compensation.

Similarly, while the demic structure of house mouse populations facilitates inbreeding, characterizing the extent of inbreeding is a complex undertaking. We model inbreeding via full brother-sister mating, which allows us to easily evaluate the impact of linking the reproductive success of relatives. The breeding structure of house mouse likely allows for other forms of related matings, notably father-daughter matings as each breeding unit usually consists of a single ‘dominant’ male (Reimer and Petras 1967). While brother-sister and father-daughter matings have the same inbreeding coefficient, fathers are nearly certain not to be homozygous for the sterile *t*-haplotype, while brothers are more likely to be infertile. As such, father-daughter inbreeding may be less favorable to the spread of lethal *t*-haplotypes than is brother-sister inbreeding. However, this effect is likely limited as fathers eventually die and are replaced by their sons. Other complex models of inbreeding in local groups of mice (e.g., cousin inbreeding) are also plausible and each could offer its own impact on the evolution of lethal *t*-haplotypes.

While inbreeding no doubt occurs in mice, disassortative mating is also possible. The MHC locus – which is thought to influence mate choice – is contained within the inversions making up the *t*-complex. Preliminary evidence suggests that mice can smell the difference between individuals differing only in their MHC alleles, but the extent to which this controls mate choice is still unclear (Yamazaki et al. 1976; Cheetham et al. 2007; Sherborne et al. 2007; Yamazaki and Beauchamp 2007; Lindholm et al. 2013). The association between the *t*-haplotype and MHC complex has been put forward as a possible mechanism for female mice to discriminate between individuals carrying the wild type, the same *t*-haplotype, or different *t*-haplotypes (Figueroa et al. 1985; Ben-Shlomo et al. 2007; Lindholm et al. 2013). A recent study found that females heterozygous for the *t*-haplotype exhibited an MHC-dependent fertilization bias in both the choice of sire and the choice of sperm (Lindholm et al. 2013). Their evidence suggests that linkage between the *t*-haplotype and MHC complex may provide a mechanism by which individuals heterozygous for the *t*-haplotype selectively mate with individuals homozygous for the wild-type. Additionally, multiple mating by females – which is not uncommon in mice – has also been shown to influence *t*-haplotype evolution and result in *t* dynamics that align with natural observations (Manser et al. 2011). Polyandry would restore the fertility of sisters who mated once with a sterile brother. As such, numerous features of mouse mating structure could temper the effect of inbreeding on the evolution of lethal *t*-haplotypes.

A few features of our model warrant additional considerations. We assumed no fitness impact of the *t*-haplotype when heterozygous and no impact of the sterile distorting *t*-haplotype on female fitness. This latter assumption is likely important for the evolutionary replacement of sterile *t*-haplotypes with lethal ones, as the extent of compensation required to make up for the death of a female with reduced fitness is obviously less than that required to make up for the death of a female with high fitness. We also assumed no explicit model of space or migration – let alone the effect of the *t*-haplotype on migration propensity. Females competing with sisters for space could have similar effects as reproductive compensation as the fitness of a litter may not decrease linearly with the number of living daughters.

Ultimately, our results provide support for an adaptive explanation of *t*-haplotype evolution; however, given the complexities of mouse population structure and the biology underlying reproductive compensation (see above), we suggest as serious consideration of non-adaptive explanations for the replacement of sterile *t*-haplotypes by lethals. Charlesworth (1994) argued that lethal *t*-haplotypes required an adaptive explanation – rather than models of drift or hitchhiking – because no such lethal haplotypes are found in SD, a similar distorter system in *Drosophila*. Certainly differences in the reproductive biology of *Drosophila* and *Mus* (e.g., mice get pregnant and nurse their pups) are consistent with a possibility that there is substantial reproductive compensation in mice and not *Drosophila*. However, other differences between these taxa may make non-adaptive explanations more plausible in mice than flies. Namely, while the evolution of lethality by drift may seem far-fetched, the low frequency of the *t*-haplotype, the linkage of MHC to the *t*-haplotype, and the small breeding unit size of mice (as compared to flies) makes selection against recessive traits on the *t*-hapltoype rare and ineffective. As such, the non-adaptive evolution of lethal *t*-haplotypes – although not formally modeled here – is a serious hypothesis which is consistent with the higher frequency of *t*-haplotypes in populations with sublethal and/or complementing lethals (i.e., two *t*-haplotypes which are lethal when homozygous, but not lethal when an individual has one of each such allele) than in populations with single lethal *t*-haplotypes (Ardlie and Silver 1998). An adaptive and non-adaptive mechanism for *t*-haplotype evolution need not be conflicting, and, instead, may reflect various routes for lethal *t*-haplotypes to invade depending on its time of action (or extent of reproductive compensation) and the structure of the host population.

## Author Contacts

Manisha Munasinghe –

Yaniv Brandvain –

## Author Contributions

M.M and Y.B conceived the study and designed all modeling framework. M.M. wrote and ran all model scripts. M.M. visualized the results and wrote the first draft. All authors contributed to the writing of the manuscript. Y.B. supervised the project.

## Acknowledgements

We would like to thank Anna Lindholm and Andrew Clark for their useful comments, insights, and suggestions regarding model development and biological context.

## Data Availability

The scripts for all simulations, job submissions, simulation results, and figure visualizations can be found on GitHub: https://github.com/mam737/tHaplotype_Inbreeding.

---

Figure S1 **Final Haplotype Frequency For Model 3 With Reproductive Compensation Without Inbreeding.** As a proof of concept, we replicated Charlesworth 1994’s model for the evolution of lethal *t*-haplotype given reproductive compensation. We show that our results match those of Charlesworth 1994 (see Figure 2 in Charlesworth 1994). The x-axis shows the extent of reproductive compensation, while the y-axis shows the equilibria frequency. The dark brown line shows the frequency of the lethal *t*-haplotype on its own, while the lighter green line shows the overall frequency of the *t*-haplotype (the lethal + male-sterile). This shows that the lethal haplotype only invades under strong amounts of reproductive compensation (C > 0.8) and that it does not displace the present male-sterile haplotype.

Figure S2 **Equilibria Frequency Comparison Between Dunn and Levene 1961 and Model 3.** Dunn and Levene 1961 characterized analytical predictions for the equilibrium frequencies for a lethal *t*-haplotype (*s* = 0, *t* = 1.0) that invades a naïve population. We consider a special case of Model 3, where *t* = 1.0, *C* = 0, and we only introduce the lethal haplotype, and show that our model results perfectly align with Dunn and Levene’s predictions. Both results show that a lethal haplotype can invade a naïve population and reach high appreciable frequency, but these values are often higher than observed estimates.

