## Supplementary material for "Together Inbreeding and Reproductive Compensation Favor Lethal *t*-Haplotypes": MM_YB_Supplement

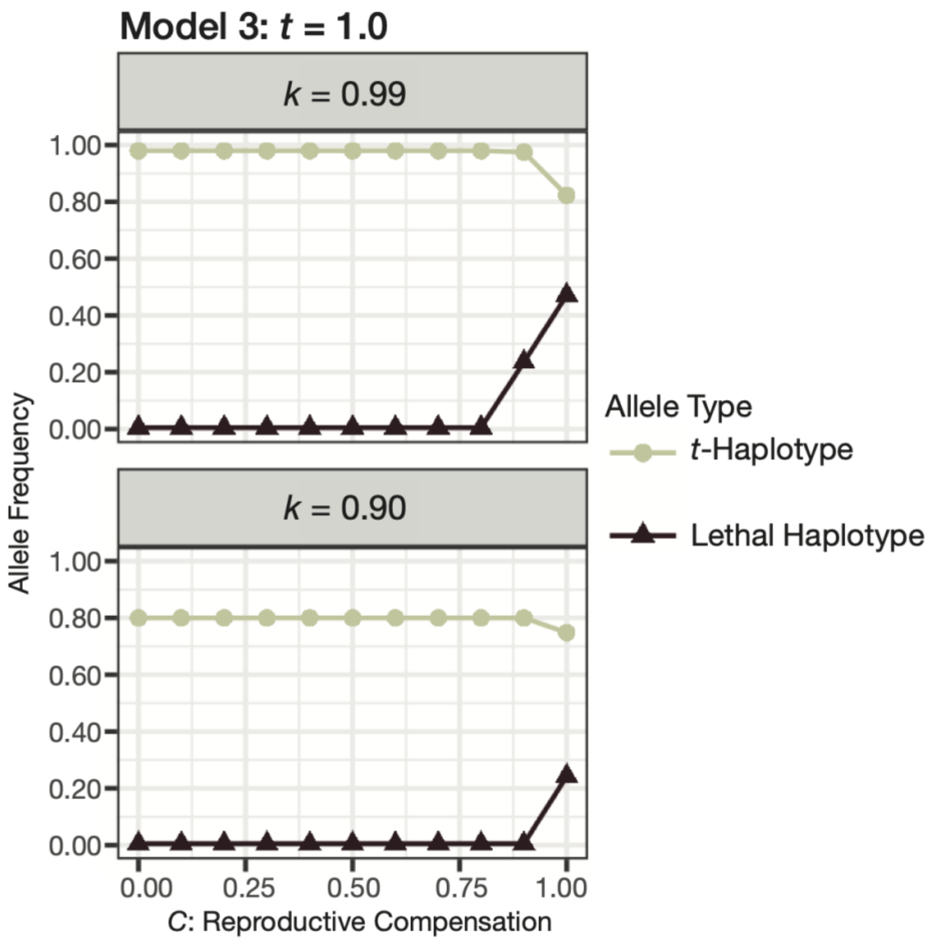


**Figure S1 Final Haplotype Frequency For Model 3 With Reproductive Compensation Without Inbreeding.** As a proof of concept, we replicated Charlesworth 1994's model for the evolution of lethal *t*-haplotype given reproductive compensation. We show that our results match those of Charlesworth 1994 (see Figure 2 in Charlesworth 1994). The x-axis shows the extent of reproductive compensation, while the y-axis shows the equilibria frequency. The dark brown line shows the frequency of the lethal *t*-haplotype on its own, while the lighter green line shows the overall frequency of the *t*-haplotype (the lethal + male-sterile). This shows that the lethal haplotype only invades under strong amounts of reproductive compensation (C > 0.8) and that it does not displace the present male-sterile haplotype.

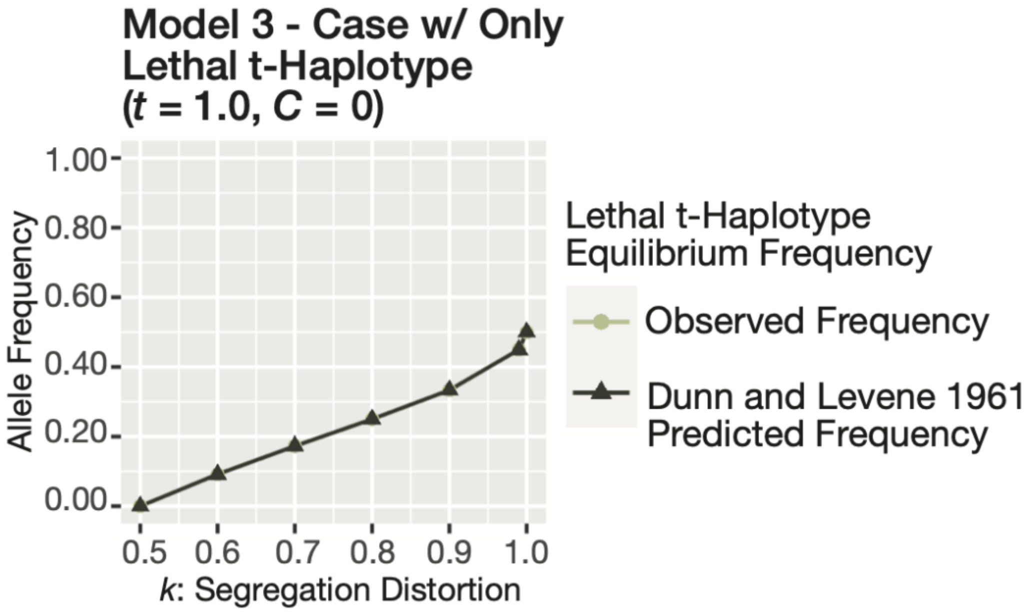


**Figure S2 Equilibria Frequency Comparison Between Dunn and Levene 1961 and Model 3.** Dunn and Levene 1961 characterized analytical predictions for the equilibrium frequencies for a lethal *t*-haplotype ($s = 0$, $t = 1.0$) that invades a naïve population. We consider a special case of Model 3, where $t = 1.0$, $C = 0$, and we only introduce the lethal haplotype, and show that our model results perfectly align with Dunn and Levene's predictions. Both results show that a lethal haplotype can invade a naïve population and reach high appreciable frequency, but these values are often higher than observed estimates.

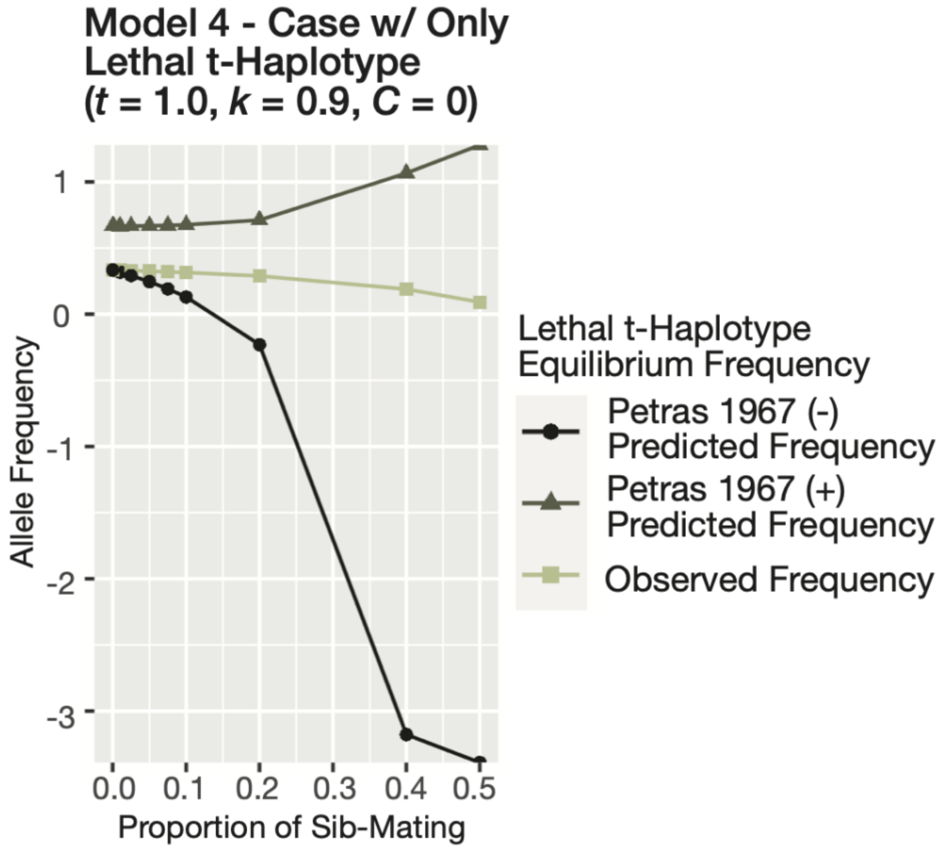


**Figure S3 Equilibria Frequency Comparison Between Petras 1967 and Model 4.** Petras 1967 characterized analytical predictions for the equilibrium frequencies of a lethal *t*-haplotype ($s$ = 0, $t$ = 1.0) that invades a naïve population. Petras 1967 incorporates inbreeding into their Model via deviations from Hardy-Weinberg equilibrium due to population structure. We could roughly compare these results to Model 4, where $t$ = 0, $k$ = 0.9, and $C$ = 0. We show that under low proportions of sib-mating our results roughly align with the results in Petras 1967. However, under higher proportions of sib-mating the comparison between our model and Petras 1967 breaks down, resulting in unreasonable predicted frequencies using the Petras 1967 expectations.
